# Correction for both common and rare cell types in blood is important to identify genes that correlate with age

**DOI:** 10.1101/2020.05.28.120600

**Authors:** Damiano Pellegrino Coppola, Annique Claringbould, Maartje Stutvoet, BIOS Consortium, Dorret I. Boomsma, M. Arfan Ikram, Eline Slagboom, Harm-Jan Westra, Lude Franke

## Abstract

**Background:** Aging is a multifactorial process that affects multiple tissues and is characterized by changes in homeostasis over time, leading to increased morbidity. Whole blood gene expression signatures have been associated with aging and have been used to gain information on its biological mechanisms, which are still not fully understood. However, blood is composed of many cell types whose proportions in blood vary with age. As a result, previously observed associations between gene expression levels and aging might be driven by cell type composition rather than intracellular aging mechanisms. To overcome this, previous aging studies already accounted for major cell types, but the possibility that the reported associations are false positives driven by less prevalent cell subtypes remains.

**Results:** Here, we compared the regression model from our previous work to an extended model that corrects for 33 additional white blood cell subtypes. Both models were applied to whole blood gene expression data from 3165 individuals belonging to the general population (age range of 18-81 years). We evaluated that the new model is a better fit for the data and it identified fewer genes associated with aging (625, compared to the 2808 of the initial model; P ≤ 2.5 × 10^−6^). Moreover, 511 genes (∼18% of the 2,808 genes identified by the initial model) were found using both models, indicating that the other previously reported genes could be proxies for less abundant cell types. In particular, functional enrichment of the genes identified by the new model highlighted pathways and GO terms specifically associated with platelet activity.

**Conclusions:** We conclude that gene expression analyses in blood strongly benefit from correction for both common and rare blood cell types, and recommend using blood-cell count estimates as standard covariates when studying whole blood gene expression.

## Background

Aging, defined as a time-dependent process characterized by physical and cognitive decline, is one of the main risk factors for autoimmune diseases, neurodegenerative diseases, cancer and diabetes [1,2]. To better understand this process on a molecular level, changes in gene expression during aging have been previously studied in whole blood [3,4]. However, blood contains many cell populations, such as white blood cells (WBC) that can be divided into granulocytes, lymphocytes and monocytes, and further into more specific WBC subtypes [5]. Since the proportions of these cell populations vary with age [6–9], it is necessary to correct for cell counts when using gene expression from blood. Indeed, uncorrected gene expression data from whole blood has been shown before to be biased by the gene expression pattern of the most abundant cell type at the moment of sampling [10].

Here, to better identify cell-independent transcriptional signatures during aging, we expanded the regression model that corrects for the number of WBC presented in our previous work [3] (hereafter called Initial Model, IM), by taking into account additional specific WBC subtype counts in our new model (hereafter called Extended Model, EM). We compared the performance of these two models in a meta-analysis using 3165 human peripheral blood-derived RNA-seq samples from four independent Dutch cohorts present in the BIOS consortium, namely LifeLines Deep, Leiden Longevity Study, Netherlands Twin Registry and Rotterdam Study [11–14]. Further, we show that the EM complies with the assumptions of linear regression and provides a better fit to the data as residuals decrease. Lastly, we analyze the genes significantly up- and downregulated by functional enrichment in order to understand to which extent the models and cell correction can be used to extract biological information regarding aging in a general population.

## Results

### Improved cell correction is necessary to identify cell-independent gene expression patterns

We performed an association of gene expression changes with age using data from four Dutch cohorts (Tab. S1). To take into account the differences in the data, we conducted a meta-analysis across these cohorts. We included only samples with all categorical covariates reported, leaving a total of 3165 individuals (Tab. S1). An overview of this study is presented in Fig. 1.

**Fig. 1.**
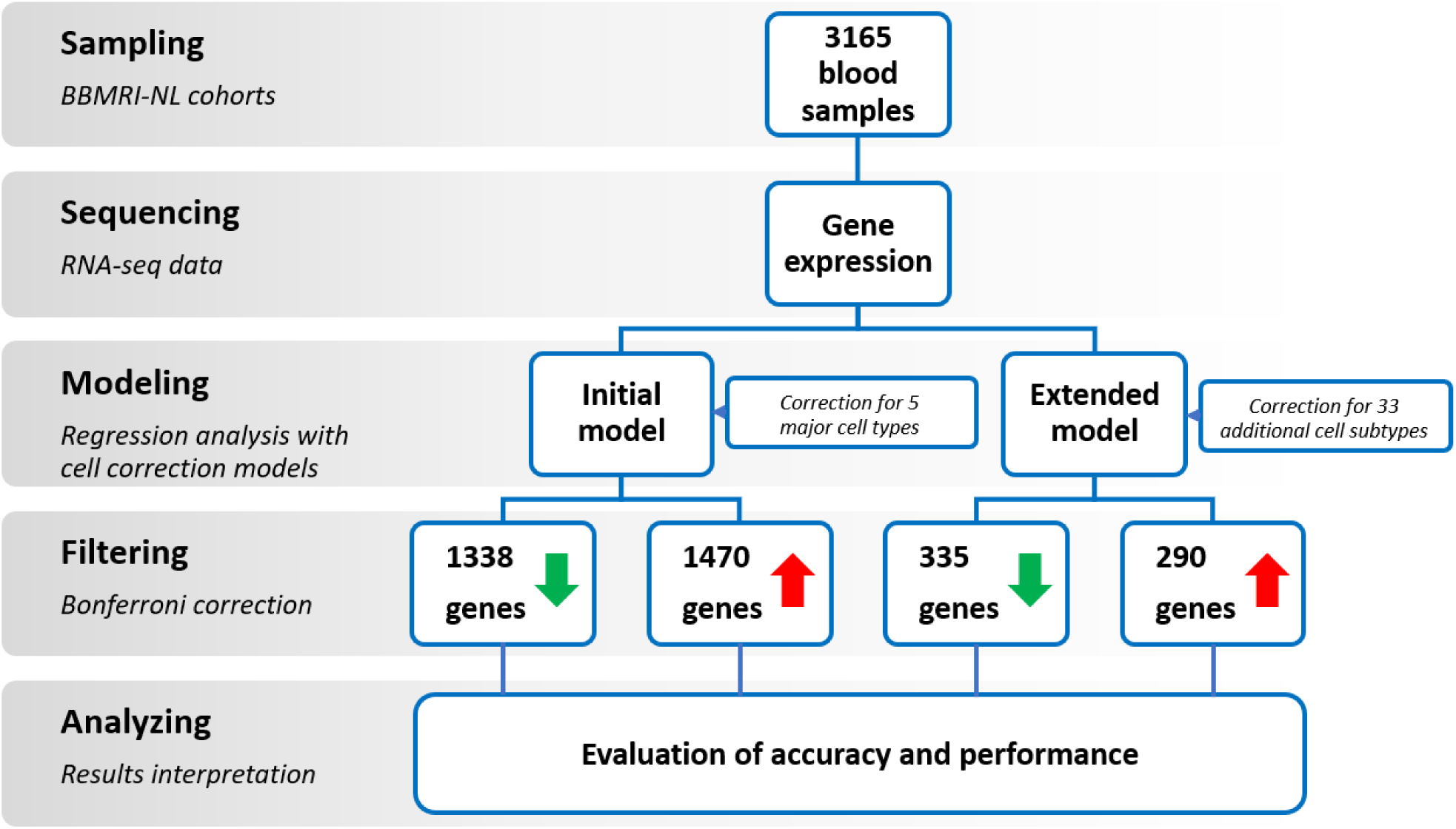
Overview of this study. 3165 complete samples from four BBMRI-NL BIOS consortium cohorts were used (see text for details). Gene expression was related to age and selected covariates depending on the regression model applied (Initial or Extended). Genes significantly associated with age were retrieved by applying Bonferroni correction (P ≤ 2.5*10^−6^) and gene lists obtained were compared to establish the efficiency of the models and analyzed to get insights on the process of aging.

We tested 19932 genes expressed in blood and analyzed the data by applying two models, the IM and the EM (see *Methods* and Fig. 1). The IM was presented previously [3]: it accounts for the main WBC types (number of granulocytes, lymphocytes, monocytes), erythrocytes and platelets, while our new model here presented, EM, corrects for 33 additional WBC subtypes (see *Methods*, Tab. S2 and S3). These additional WBC subtypes were imputed with Decon-cell [15]. We observed small but significant correlations between age and most measured and imputed cell counts (Fig. S1), presenting evidence that adding WBC subtypes is beneficial for the correction models. For example, different imputed cell types, such as naïve CD8^+^ subtypes (IT50 and IT54 [16]), show a strong negative correlation with age when considering both the overall (Fig. S1) and the single cohorts (data not shown).

Using the IM, we identified 1338 genes significantly downregulated and 1470 upregulated with age after Bonferroni correction (P ≤ 2.5 × 10^−6^) (Tab. S4 and Fig. 1). The EM, however, reduced the number of results substantially: we identified 335 downregulated and 290 upregulated genes significantly associated with aging at the same significance threshold (Tab. S4 and Fig. 1). This decrease was expected, as many of the results from the IM may have been driven by the composition of less prominent cell types that were included in our EM model. While 511 out of 625 EM genes were also present in the IM results, the 114 additional EM genes were only detected after rigorous correction for cell types (Fig. S2). To validate our results, we compared the number of genes retrieved through our models with the 1497 genes reported in our previous work [3] (gene set 1, GS1) and the 481 genes identified by Lin and colleagues [4] (gene set 2, GS2), a study that uses a slightly different correction model to study aging. As reported in Tab. S5, the highest number of overlapping genes was found between the IM and the GS1 (672, 24% of our 2808 IM genes). Considering that the number of tested genes is different (11908 for GS1 and 19932 for IM, 10890 in common), this overlap is quite large. Moreover, all genes had the same direction of association with age. These results are unsurprising, because we used the same previous correction model [3]. When comparing the EM results with the GS1, the number of overlapping genes decreased (172, 28% of our EM genes) but the majority still had the same direction (98%). The lowest number of overlapping genes was found between the EM results and the GS2 (9 genes overlapping, 7 with the same direction). In general, differences in the number of overlapping genes may result from: 1) differences in the model used, 2) differences in the technical analyses performed [17] and 3) differences between the genes used in the discovery phase. Overall, the models show a good conservation of direction for overlapping genes, which indicates that correcting for cell populations identifies common whole blood gene expression patterns.

### The Extended Model performs better than the Initial Model

We next investigated whether both IM and EM met assumptions of linear regression. To this end, we analyzed the mean squared errors (MSE), the distribution of gene expression residuals and their homoscedasticity after applying the IM and EM. We first analyzed the impact of adding additional terms to our regression models on the MSE. As expected, MSE values of the regressions for every gene decreased when applying the EM (total EM median MSE value: 0.267, total IM median MSE value: 0.334) (Fig. 2A-B and Tab. S6). We next created QQ-plots and calculated the Pearson correlation coefficient between the observed and expected distributions to assess normality. For most genes, including the 511 shared between IM and EM, we found that applying the EM resulted in more normally distributed residual values and the correlation values were higher (total EM median r value: 0.995, total IM median r value: 0.994) (Fig. 2C-D, Tab. S6 and S7). Lastly, we wanted to evaluate heteroskedasticity (i.e. the skewness on the distribution of residuals), as this can indicate a relation between the error and the explained variable, violating the model assumptions. For this purpose, we created a modified version of both models that included all covariates with the exception of age and applied the four resulting models (IM, EM, IM-age, EM-age) in each cohort. Then, we used the rank-based Spearman correlations to correlate gene expression residuals with age [18,19]. We checked the normality of these Spearman ρ values and meta-analyzed them across the cohorts (Fig. S3). We observed that the absolute correlations were smallest in the EM model (EM median value: 9 × 10^−3^), and largest in the IM without age (IM-age median value: 6 × 10^−2^) (Fig. 2E-F, Tab. S6 and S7). Large ρ values indicate a less precise prediction and larger errors. In general, the EM performs better than the IM, and it is specifically noteworthy that the EM without age performs better than the IM without age. Adding cell counts clearly improves the prediction of gene expression values. These three analyses indicate that the EM satisfies the assumptions of linear regression better than IM. Moreover, adding cell counts as covariates improves reliable identification of aging-related genes in whole blood.

**Fig. 2.**
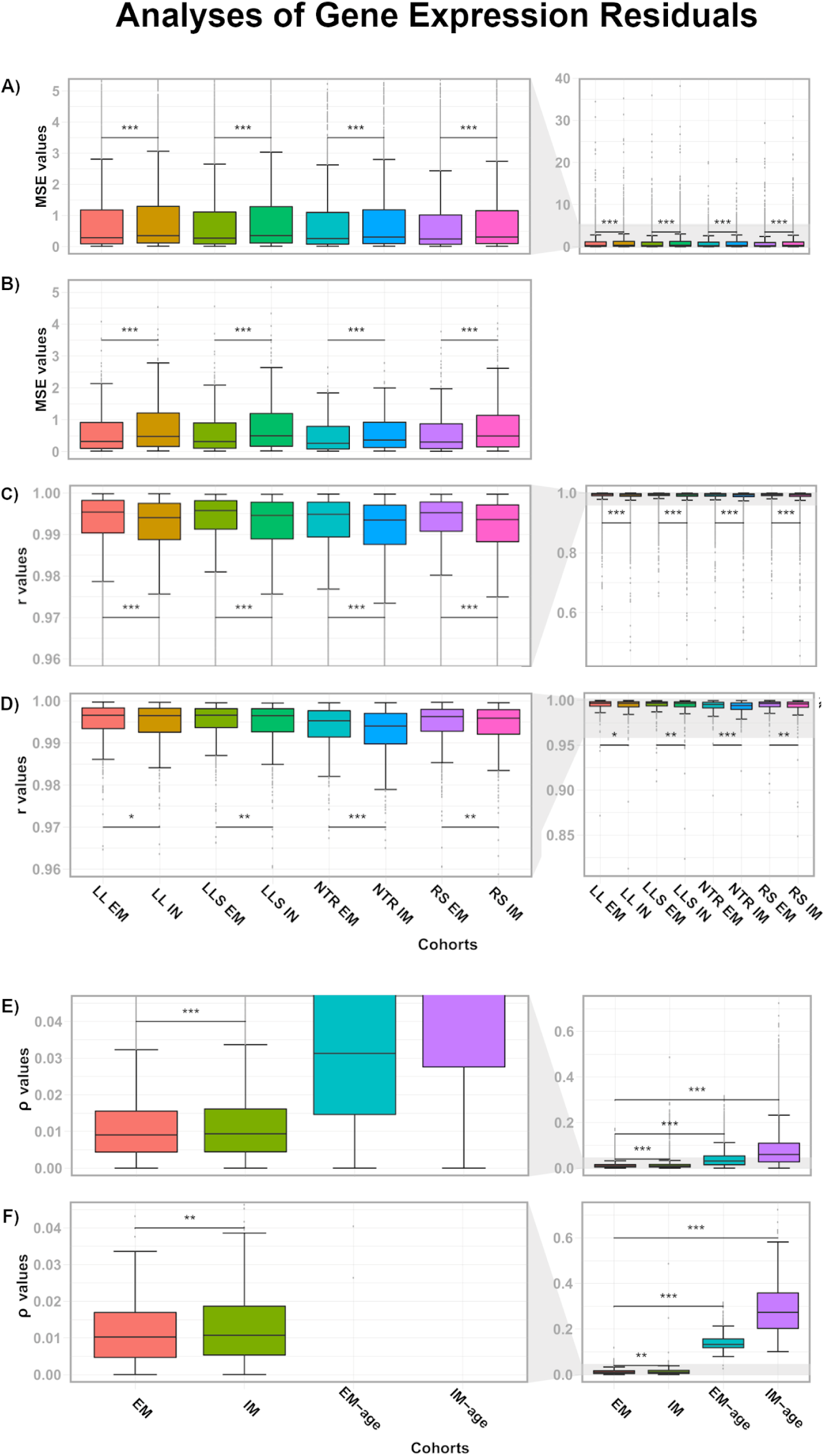
Gene expression residuals decrease with the EM. MSE values for regressions related to genes in every cohort after applying the IM and the EM are reported for all genes in (A), and the 511 shared genes in (B). QQ plot Pearson correlation coefficients (r values) related to the distributions of gene expression residuals are shown for all genes in (C) and for the shared genes significantly associated to aging in (D), after applying the IM and EM models. Homoscedasticity was evaluated by correlating gene expression residuals from every model with age, and the absolute Spearman ρ values obtained after meta-analysis are reported for all genes (E) and the shared genes significantly associated with aging (F). LL, LifeLines DEEP; LLS, Leiden Longevity Study; NTR, Netherlands Twin Registry; RS, Rotterdam Study; EM, extended model; IM, initial model; EM-age, extended model without age as covariate; IM-age, initial model without age as covariate. Statistical significance was assessed with a paired, one-tailed Wilcoxon test. The stars indicate statistical significance: *** P ≤ 0.001, ** P ≤ 0.01, * P ≤ 0.05.

### Single-cell RNA-seq data reveals the contribution of cell types to gene expression during aging

Every cell type has its own gene expression pattern, so the composition of blood cells influences the total gene expression observed in whole blood RNA-seq data. To test to which extent the aging-related genes found by the models were influenced by blood cell populations, we investigated the mean expression of these aging-related genes in single-cell RNA-seq (scRNA-seq) data of 11 different blood cell types [20]. As shown in the t-SNE plots (Fig. 3A), aging-related genes retrieved through the IM have a propensity to be expressed in specific parts of the t-SNE plot that match with cell types, while EM genes maintain a lower and more stable expression across cell types (Wilcoxon test, P ≤ 2.2 × 10^−16^, Fig. 3B on the left), suggesting that it is not a specific cell type driving the associations. Secondly, we used differential expression patterns to identify blood cell type specific markers in the list of IM or EM significant aging-related genes, and visualized the mean expression in t-SNE plots (Fig. S4). The EM aging-related genes contain fewer cell type specific markers: no markers could be identified for three cell types (Natural Killer bright subset, CD8^+^ T and B cells). Importantly, the cell type marker genes that were identified among EM genes are less representative for their cell types than the IM markers, as shown in Fig. S4. In addition, we observed that the mean expression range for the EM genes was always larger, highlighting a higher gene expression variation (mean expression of IM genes per cell: 0.05 – 0.21; EM: 0.02 – 0.28, Fig. 3A and S4). This observation was supported by the scRNA-seq coefficients of variation calculated for the EM genes (Wilcoxon test, P = 3.794 × 10^−10^, Fig. 3B on the right). In summary, the scRNA-seq data indicate that EM genes are less driven by cell types than the IM genes, suggesting that the EM model enables a better identification in blood of cell-quantity independent genes related to aging.

**Fig. 3.**
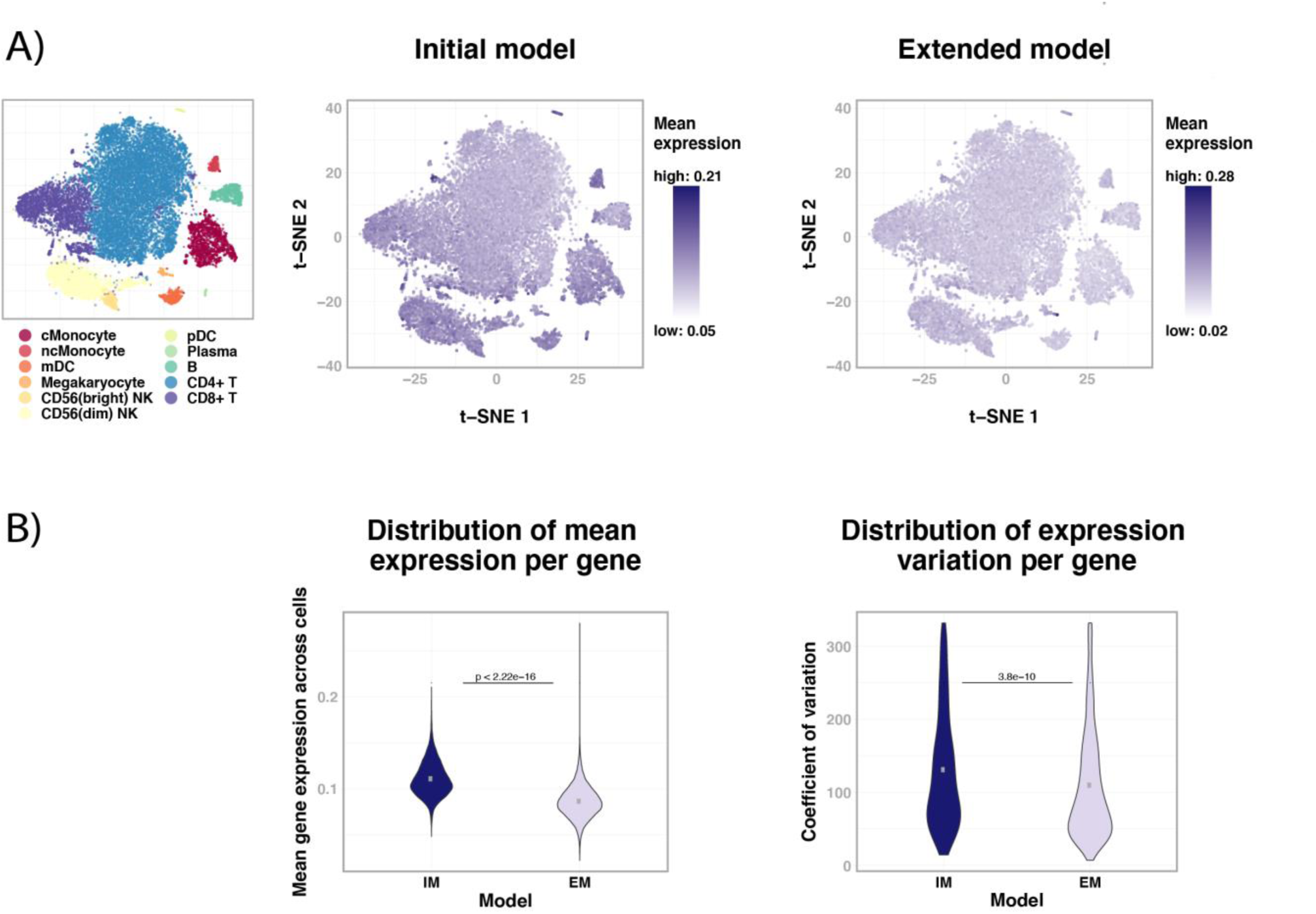
scRNA-seq data reveals that the aging-related genes found with the EM are not related to specific blood cell populations. The mean expression value of aging-related genes is plotted in relation to blood cell populations in A): the upper t-SNE plots refer to the IM, on the left, and the EM, on the right. In B), the distributions of the means for gene expression for every cell in the above t-SNE plots according to the IM and EM are reported on the left, while on the right the distributions of the coefficient of variation are presented for both the IM and EM. Statistical significance was assessed with a Wilcoxon test. The level of statistical significance was set at P ≤ 0.05. For details regarding cell population-specific regions, refer to [20].

### Functional enrichment analysis and aging signatures

In order to investigate whether the EM-derived aging-related genes were more informative than the IM-derived genes, we performed functional enrichment using Enrichr [21]. As 82% of the EM genes were also present in the IM list, we expected comparable functional enrichments. On the contrary, very few pathways and GO terms were shared between the EM and IM lists (Tab. S8). The fact that we observed a smaller number of genes in the EM list did not translate to a lower number of EM-specific enrichments. Therefore, we hypothesized that although a high number of genes is shared between the EM and the IM, the difference in the functional enrichment results was due to the exclusion of genes that are influenced by cell quantity, for which the IM did not correct. Indeed, the enrichments for the EM genes clustered around potential aging-related mechanisms. For example, changes in GO biological processes ascribable to the regulation of gene expression were downregulated (e.g. ‘regulation of transcription, DNA-templated’ – GO:0006355, ‘regulation of nucleic acid-templated transcription’ – GO:1903506, ‘regulation of protein processing’ – GO:0070613), in agreement with previous findings [3] and the IM results.

Hemostasis, the process to prevent and stop bleeding, emerged as a key upregulated pathway from the various EM-related enrichment analyses (Tab. S8). The Kegg pathway ‘coagulation cascade’ and the Reactome pathway ‘hemostasis’ were both significantly upregulated (P ≤ 6.2 × 10^−4^, P ≤ 4.5 × 10^−6^, respectively), suggesting that changes in the expression of genes related to hemostasis and platelet functioning during aging have a very robust signature, as previously reported [22–26]. Changes in GO biological process terms related to platelet activity (GO:0045055, GO:0002576, Tab. S8) and GO cellular compartment terms linked to platelet granules (e.g. ‘platelet alpha granule’ – GO:0031091, Tab. S8) were also found to be significant. Notably, both models included the correction for platelet counts, suggesting that these functional enrichments described the activity of platelets independently of their prevalence. Platelet count remained more or less stable during aging in our data (Fig. S1), so the number of platelets is not expected to drive these enrichments.

After applying the EM, we expected that genes involved in the same biological process and under the same regulation could show a common pattern. To identify this pattern, we calculated the correlations between the gene expression residuals. We observed several clusters with highly correlating values (Fig. 4 and Fig. S5), which we further analyzed with Enrichr. While most clusters did not show a clear enrichment, cluster 1 of the upregulated EM aging-related genes (Fig. 4, upper left corner) was enriched for terms related to platelet activity, again highlighting its role in aging. Five genes (*PF4, PPBP, STON2, MYLK, LMNA*) from the platelet-related cluster 1 were previously identified to be differentially expressed with age in platelets [25]. Although *PF4* and *PPBP* did not show the same direction of effect, a difference that may result from the sample size or the model used, the overall finding that platelets show increased activity with age is conserved [22,23,25,26].

**Fig. 4.**
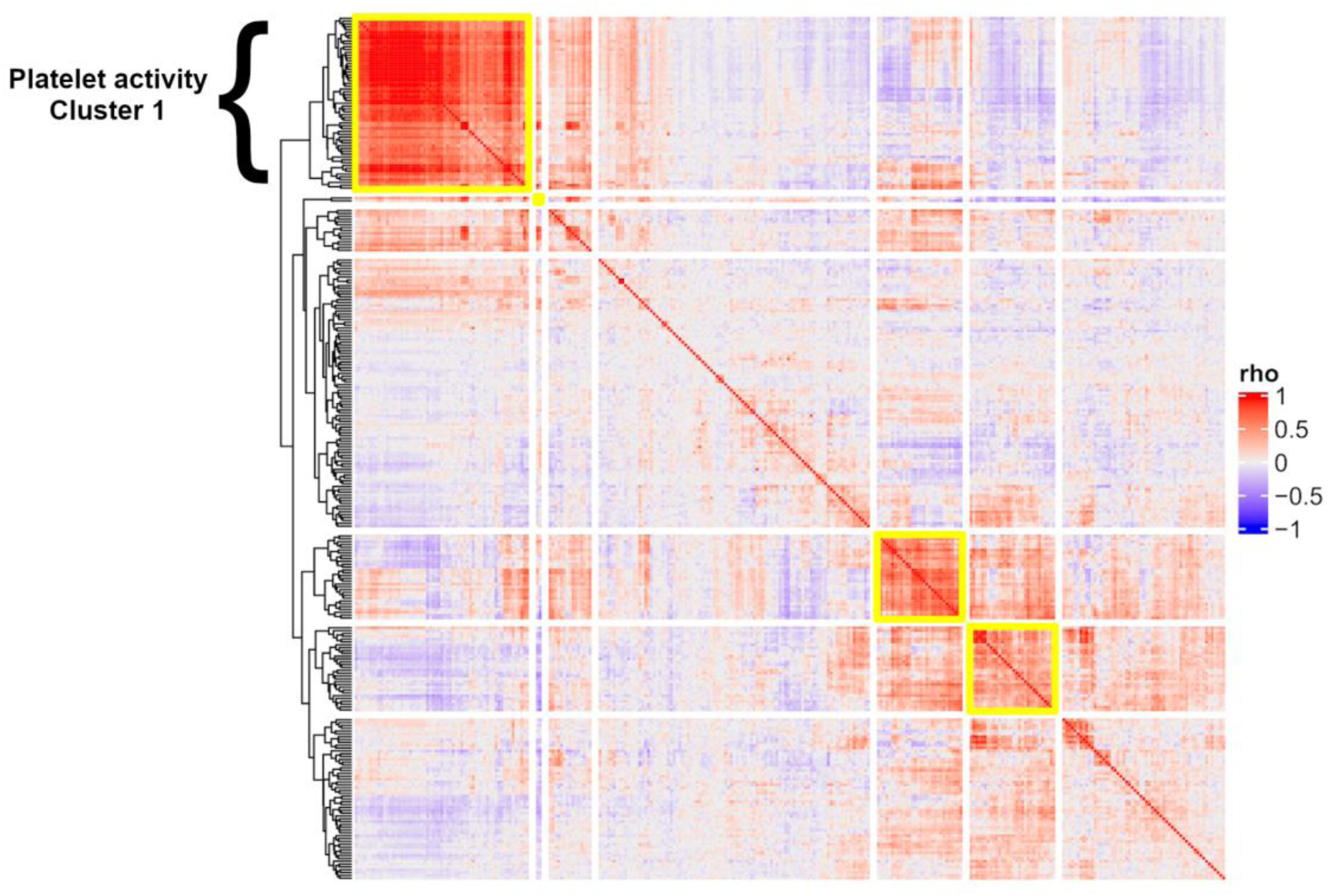
Heatmap of gene expression residuals correlations for EM upregulated aging-related genes. Upregulated EM aging-related genes were clustered based on the correlations of gene expression residuals and highly correlating clusters were identified and highlighted with a yellow border. The cluster in the upper left corner contains genes associated with platelet activity pathway and GO terms.

## Discussion

Aging is a process that enhances the probability of getting diseases such as cancer, diabetes and various types of neurodegenerations. In order to understand how an organism reaches these diseased states, it is valuable to study the preceding period, where the organism ages. Changes can be investigated by analyzing aging cohorts as representatives of an aging population. Following this reasoning, in this study we used four Dutch aging cohorts (Tab. S1) and analyzed gene expression changes during aging in whole blood, an easily accessible tissue, by implementing a new model (EM) to correct for cell type proportions. This extended cell correction enabled us to calibrate gene expression according to the number of blood cells and extract an aging gene expression pattern that was less influenced by cell quantity compared to previously published models [3,4]. To test the performance of our EM, we evaluated its compliance to the assumptions of regression. The EM outperformed the old model, IM, when analyzing the MSE, normality of residuals and homoscedasticity, highlighting that an increased cell correction results in a more accurate gene expression estimation during aging.

Next, we asked which cell population contributed the most to the list of aging-related genes provided by both the IM and EM. For this purpose, we calculated per cell type the mean gene expression of both IM and EM genes using scRNA-seq data from ∼25000 blood mononuclear cells of 45 donors [20]. The EM aging-related genes had lower mean gene expression levels, fewer cell type specific marker genes and those markers that were present were less abundantly expressed (Fig. S4). We consequently reasoned that these genes are less influenced by cell composition and quantity.

We performed a functional enrichment analysis for GO terms, Kegg and Reactome pathways in order to gain insight on the blood-based biological mechanisms driving aging. Although many of the EM genes were also identified using the IM, the enrichments were often not overlapping suggesting an increased precision in evaluating the relation between gene expression and age. In particular, platelet-related categories stood out in these results. We clustered the EM genes based on gene expression residuals and again found the strongest enrichment in the upregulation of platelet activity.

Since our EM includes a correction for platelet counts, the observation that platelet activation is enriched in relation to the EM aging-related genes is possibly due to the following reasons: 1) the EM did not correct for cell counts sufficiently or 2) an increase in platelet activity is a true signature of aging. While we cannot exclude the first reason, the fact that platelets do not associate with age in our data make it less plausible. Moreover, platelet activity has been reported to increase with age in literature [22,23,25,26] and incubating human platelets with media from senescent human fibroblasts increases platelet activation and degranulation [24]. Upon degranulation, platelets release the factors present in their granules into the surrounding environment. Of note, our functional enrichment analysis retrieved GO terms related to alpha granules, which store PPBP and PF4. These proteins are known to be increasingly secreted during aging [22,27,28]. The genes encoding these proteins were found to be upregulated aging-related genes and, more specifically, they contributed to the enrichment of alpha-granule-related cell compartment GO terms (Tab. S8). Interestingly, an earlier study that performed RNA-seq within isolated platelets has observed decreased expression of *PF4* and *PPBP* with age (n = 154 [25]), while studies in whole blood show upregulation with age (current study: both genes significant; in the previous study [3]: *PF4* not tested, *PPBP* nominally significant). Within our scRNA-seq data, both genes are specifically expressed in megakaryocytes, the precursors of platelets (Fig. S6), suggesting that the observed upregulation is not driven by the expression in any other blood cell types, but by platelets or megakaryocytes themselves. Although these results may arise from differences in sample sizes or models used, this observation coupled with the fact that older individuals have higher levels of PF4 and PPBP protein in their plasma indicates that platelets become more active with age as reflected both in gene expression levels and protein abundance.

In addition, alpha granules are known to store aging-related proteins, such as IGF1, a protein that has been extensively connected to aging together with its orthologs in multiple organisms [29,30]. Therefore, an enhanced platelet degranulation itself could have a major impact on the progression of aging. In summary, we hypothesize that the platelet enrichment observed in the EM aging-related genes represents one of the molecular signatures of aging. The increased platelet activation and subsequent release of aging factors could affect other cells and in turn the whole organism. However, many details regarding the mechanisms that are affected by these aging factors remain to be discovered.

## Conclusions

Overall, we have shown that an extensive correction for cell type differences can dramatically alter the effect sizes and significance of associations between genes and age. On top of this correction for measured or imputed cell counts, we believe that large scRNA-seq datasets (e.g. sc-eQTLGen consortium [31], The Human Cell Atlas [32]) will be essential to visualize and quantify to what extent associations are independent of cell type composition. Our and previous findings [25] indicate that it will be essential to investigate to what extent the increased platelet activity is driven by megakaryocytes using larger blood-based scRNA-seq datasets [33]. Lastly, while the current study was performed in blood, other tissues also feature cell type heterogeneity. As such, we conclude that rigorous correction for cell type counts is important for studies in whole blood, and will help to better understand immune aging and other gene expression association studies.

## Methods

### Study populations

We performed a meta-analysis using 3165 human peripheral blood samples obtained from four independent Dutch cohorts: LifeLines DEEP (LLD, n = 1100) [11], Leiden Longevity Study (LLS, n = 585) [12], Netherlands Twin Registry (NTR, n = 852) [13] and Rotterdam Study (RS, n = 628) [14] with participants from a wide age range (Tab. S1). All cohorts followed similar protocols for genotyping and gene expression as part of the BIOS Consortium, an initiative of the Biobanking and Biomolecular Resources Research Infrastructure – The Netherlands [34].

### Gene expression

Gene expression data was obtained using the same protocol across all studies, as previously described [35]. Briefly, RNA was extracted from whole blood using PAXgene Blood miRNA Kit (Qiagen, California, USA) and paired-end sequenced with the Illumina HiSeq 2000 platform. After quality control by FastQC, adapters were removed and read quality trimming steps executed. Reads were aligned with STAR using GRCh37 as a reference while masking common (MAF > 1%) SNPs in the Genome of the Netherlands (GoNL) [36]. Reads were assigned to genes with HTseq using gene definitions from Ensembl v71. Subsequently, expression values for all exons of each gene were added up to represent gene expression, measured in base count per gene. Prior to normalization, population outliers were removed based on a plot of the first two principal components (PCs), calculated on non-imputed genotypes. The first step in the normalization procedure was the application of the trimmed mean of M-values normalization method [37]. Next, we removed genes with no variance, log_2_ transformed the expression matrix and Z-transformed by centering and scaling of the genes, following a previously published protocol described in detail in the online cookbook [38].

### Cell count imputation

We then performed imputation of cell counts, since these were not present for all included samples. For imputation, we only considered samples where all categorical covariates (sex, smoking status, fasting before blood sampling, RNA plate) were available (see Tab. S2 for missing values). We estimated the 33 WBC subtypes included in the EM using the R package Decon-cell, a method that quantifies cell types using expression of marker genes (Tab. S3) [15]. The red blood cell (RBC) count was imputed using multivariate imputation by chained equations (MICE) from the R package MICE version 2.30 [39] because this cannot be imputed based on gene expression values, but rather relies on the other cell type counts and other phenotypes (Tab. S2). In MICE, we used predictive mean matching, as it has the advantage of imputing missing values within the observed spectrum after creating a normal distribution [39,40]. Values outside the range of ± 3 standard deviations from the mean were removed after log_2_ transformation.

### Models for differential expression during chronological age

The IM was taken from the previous work [3] and is:

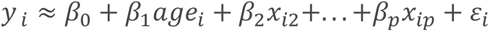

with y being gene expression levels for every gene, *i* the number of cohort samples, age (*x*_*i*1_) in years at time of blood sampling, and the following additional variables being the other covariates, including cell counts (for a total of *p* predictors). To prevent overfitting, we required at least 10 samples for each available gene [41]. As covariates, we included sex, smoking status, fasting before blood sampling, RNA plate and GC content (an RNA-sequencing quality control score). All covariates were fixed effects, except for RNA plate, which was set as a random effect. As cell counts, we included the number of RBCs, platelets, granulocytes, lymphocytes and monocytes (Tab. S1). In our EM, the imputed proportions of 33 WBC subtypes were included as additional cell count measures, to increase the power to detect cell-independent age effects. For a complete overview of WBC subtypes see Tab. S3. Both the IM and EM were tested on 19932 genes that showed expression in blood of at least 0.5 counts per million in at least 1% of the samples [42]. For these tests, we used the lmer function from the R package lme4 version 1.1.13 [43]. Sample sizes, effect directions, and P-values were extracted from the result files of both linear models.

### Meta-analysis

To combine associations across the four cohorts and to avoid bias of results due to cohort-specific effects, we first analyzed each cohort separately and then conducted a meta-analysis. We used the meta-analysis tool for genome-wide association scans (METAL) to calculate weighted Z-scores and P-values for every gene [44]. Although originally developed for meta-analysis of genome wide association studies (GWAS), METAL was easily adapted for expression associations as described in the previous work [3].

### Evaluation of the regression models

To evaluate the performance of the regression models, we used gene expression residuals and investigated MSE values, distribution of residuals and homoscedasticity. The distribution of residuals was evaluated by calculating the QQ plot Pearson correlation coefficient from sample and theoretical quantiles, considering that the higher the correlation value, the more the distribution approximates normality. Regarding homoscedasticity, meta-analysis was conducted on cohort-related, gene-specific Spearman ρ values (rho values) obtained by correlating age with the gene expression residuals, calculated from the application of the IM and EM. For this purpose, a Fisher Z-transformation was applied to the ρ values after evaluating the approximation of their distribution to normality with a QQ plot. Then, Z-scores were combined across the cohorts using a weighted approach as described in [45] and the overall Z-score converted to ρ with the inverse Fisher transformation.

### Functional enrichment analysis

To better understand gene function, we performed functional enrichment using Enrichr [21]. For this analysis, we grouped genes significantly associated with aging in either the IM or the EM into up- and downregulated genes. Using this approach, we retrieved information regarding enrichment in pathways based on KEGG and Reactome or GO terms.

### single-cell RNA-seq data and visualization

To interpret the cell type specificity of our age-associated genes, we used scRNA-seq data for approximately ∼25000 peripheral blood mononuclear cells from 45 LLD donors. Collection and normalization of the data has been described previously [20]. We used the R package Seurat version 1.4.0.13 for scRNA-seq analyses and visualizations [46]. ScRNA-seq data enabled the detection of eleven cell types: classical and non-classical monocytes, myeloid and plasmacytoid dendritic cell, CD56 bright and dim natural killer cells, CD4^+^ and CD8^+^ T-cells, B-cells, plasma cells and megakaryocytes [20]. Within these cell types, we calculated the mean expression of the genes significantly associated with aging identified by the IM and the EM, and represented their expression in t-SNE plots. We then identified genes that we considered markers for each of the 11 cell types using the function ‘FindMarkers()’ from Seurat using the loose thresholds of min.pct = 0.5, min.diff.pct = 0.2 to evaluate whether the aging-related genes were reflecting specific cell types.

## Supporting information

BIOS author information

## Declarations

### Ethics approval and consent to participate

Written informed consent was obtained previously for each of the biobanks separately in accordance with the ethical and institutional regulations [11–14].

### Consent for publication

Not applicable.

### Availability of data and materials

The data that support the findings of this study are available from BBMRI-NL but restrictions apply to the availability of these data, which were used under license for the current study, and so are not publicly available. Data are available upon reasonable request and with permission of BBMRI-NL. Summary statistics on the whole-blood gene expression, cell count imputation and expression-age associations are available from the BBMRI-NL atlas (http://bbmri.researchlumc.nl/atlas/). Raw RNA-seq data can be obtained from the European Genome-phenome Archive (EGA; accession EGAS00001001077). Individual-level genotypes are not publicly available to ensure participant privacy, but access can be requested from the BIOS consortium (https://www.bbmri.nl/acquisition-use-analyze/bios).

For the scRNA-seq data, please refer to [20].

### Authors contribution

DPC and AC contributed equally to this work.

## Acknowledgements

We thank the UMCG Genomics Coordination Center, MOLGENIS team, the UG Center for Information Technology, the UMCG research IT program and their sponsors in particular BBMRI-NL for data storage, high performance compute and web hosting infrastructure. BBMRI-NL is a research infrastructure financed by the Netherlands Organization for Scientific Research (NWO) [grant number 184.033.111].

