## Supplementary figures for "Correction for both common and rare cell types in blood is important to identify genes that correlate with age"

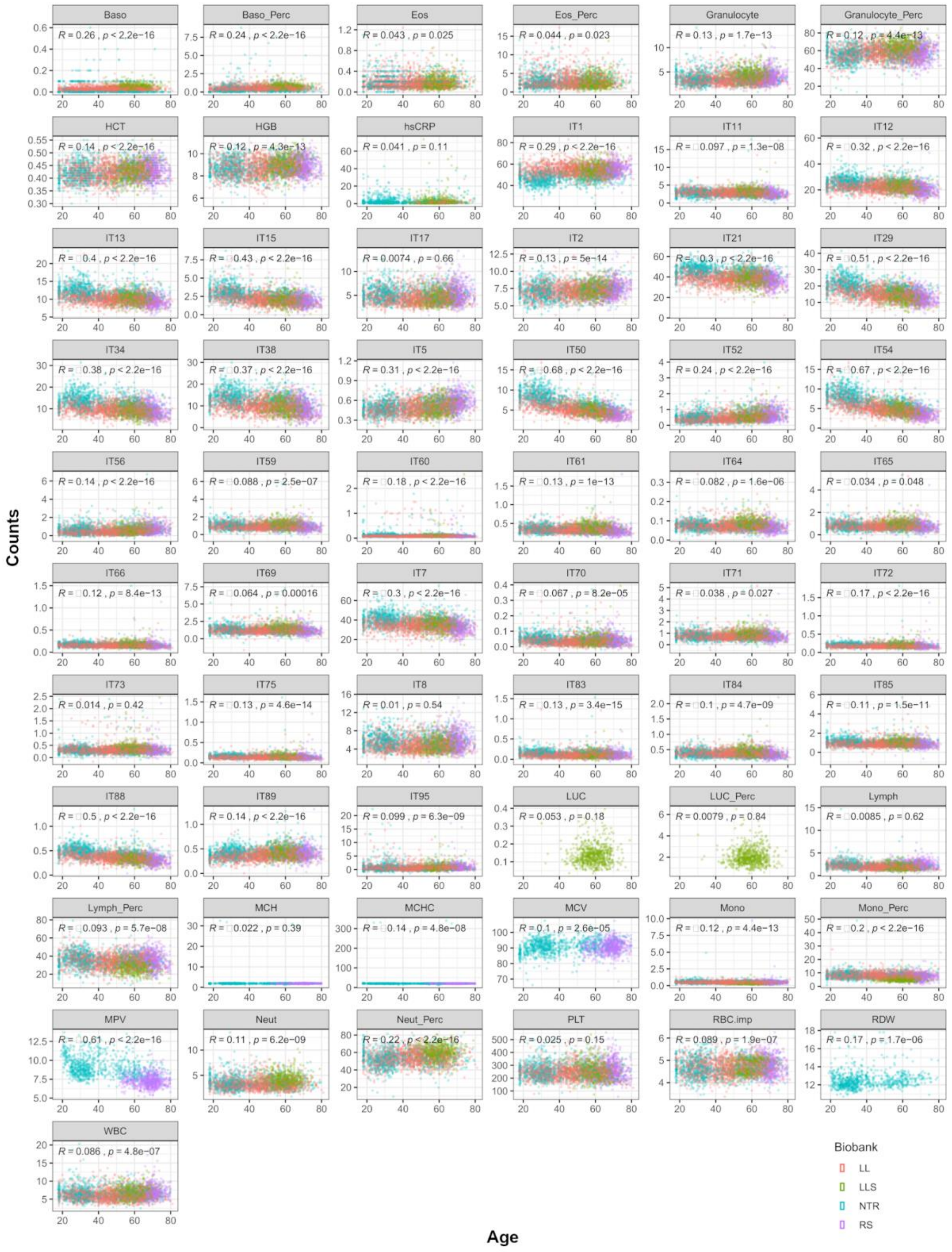


**Fig. S1. Correlations of selected variable counts with age.** The Spearman correlations of selected variables - including measured or imputed cell counts - with age are presented, colored per cohort (see legend). Baso, basophil count; baso_perc, basophil percentage; Eos, eosinophil count; Eos_perc, eosinophil percentage; Granulocyte, granulocyte count; Granulocyte_perc, granulocyte percentage; HCT, hematocrit; HGB, hemoglobin; hsCRP, high sensitivity C-reactive protein; LUC, large unstained cell count; LUC_perc, large unstained cell percentage; Lymph, lymphocyte count; Lymph_perc, lymphocyte percentage; MCH, mean corpuscular hemoglobin; MCHC, mean corpuscular hemoglobin concentration; MCV, mean corpuscular volume; Mono, monocyte count; Mono_perc, monocyte percentage; MPV, mean platelet volume; Neut, neutrophil; Neut_perc, neutrophil percentage; PLT, platelets; RBC, red blood cells; RDW, red cell distribution width; WBC, total white blood cell count; all the IT are imputed cell counts, see Table S3 for the list of names. See *Results* section for details.

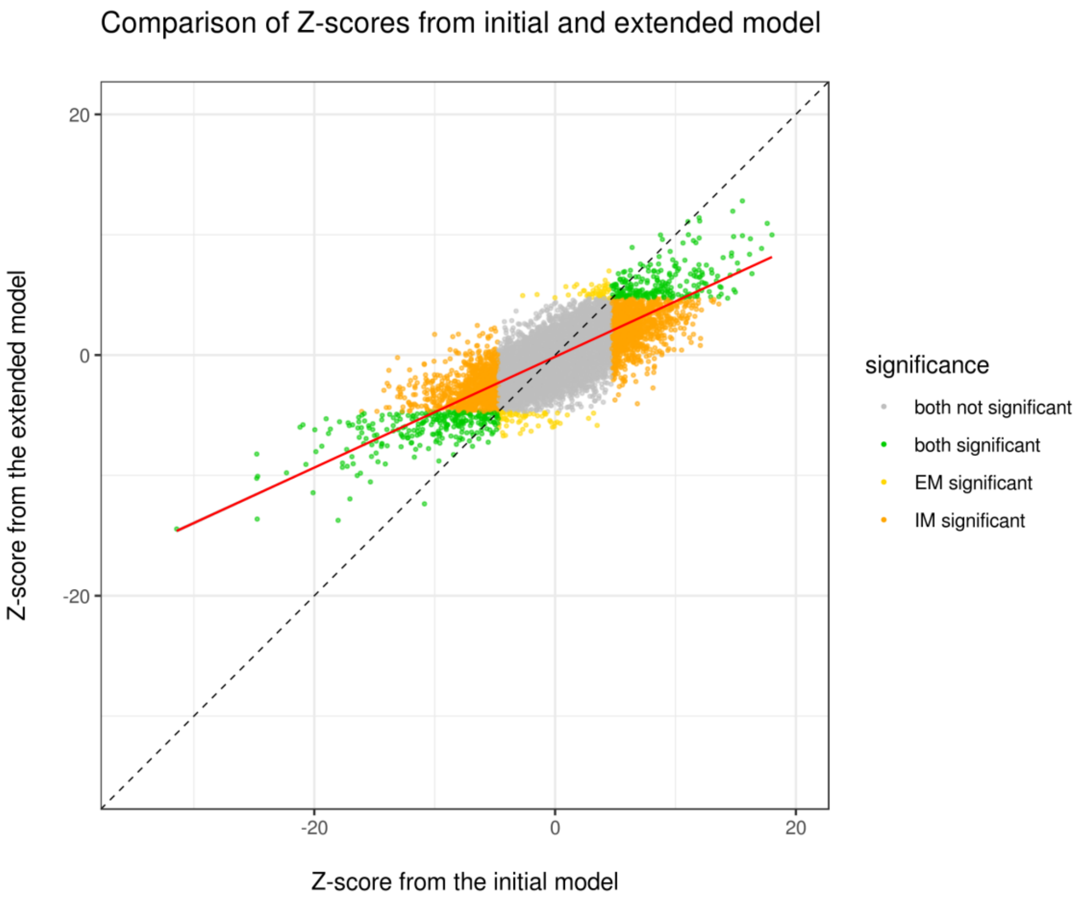


**Fig. S2. Correlation of Z-scores associated with IM and EM genes.** A Pearson correlation of the Z-scores associated with both significant and not significant IM and EM genes is shown. The 45° diagonal is presented as dashed, the correlation line is in red. See *Results* section for details.

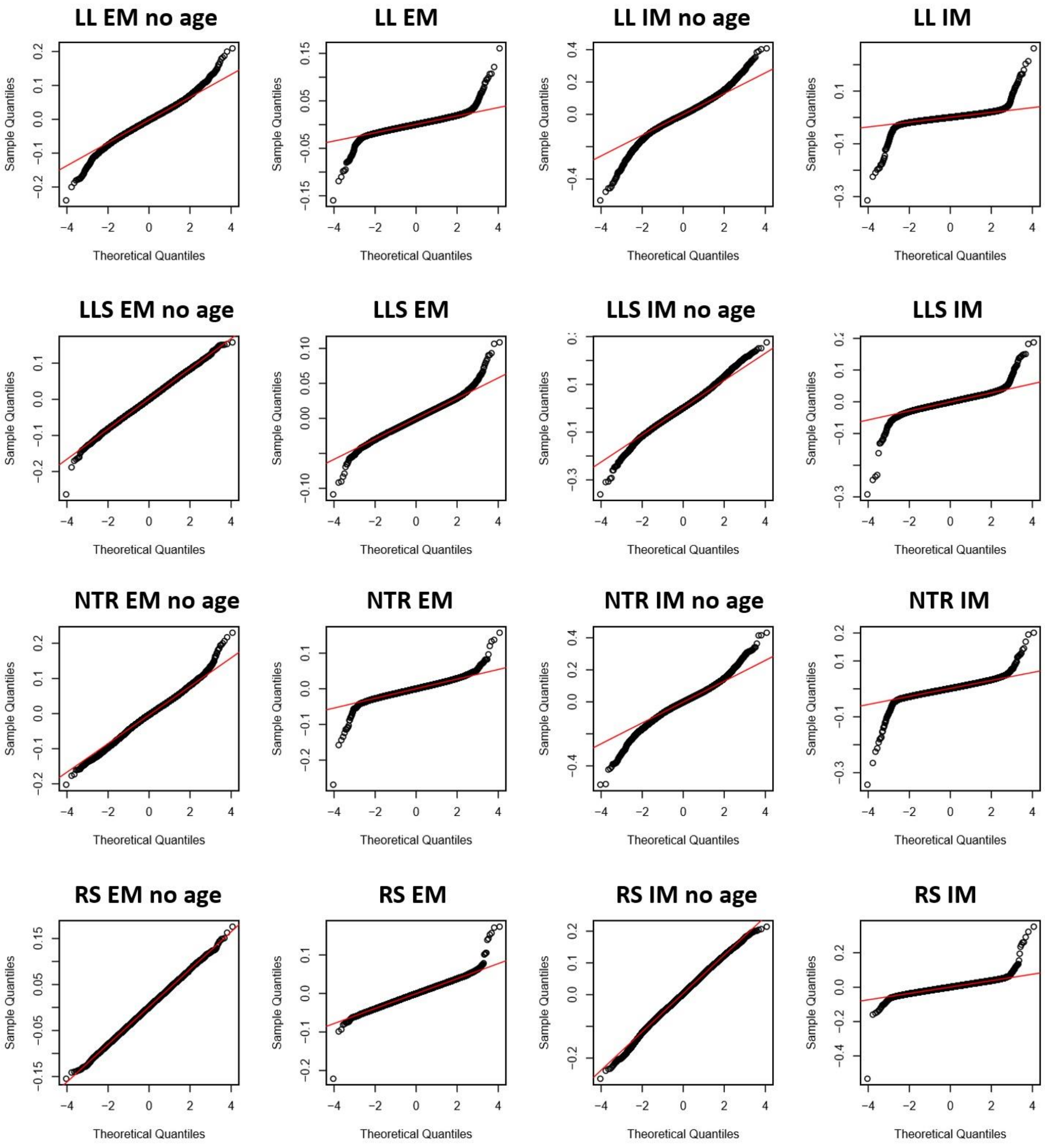


**Fig. S3. QQ plots of cohort-related, gene-specific ρ values.** QQ plots used to evaluate the distribution pattern of cohort-related, gene-specific ρ values. LL, LifeLines DEEP; LLS, Leiden Longevity Study; NTR, Netherlands Twin Registry; RS, Rotterdam Study; EM, extended model; IM, initial model; EM no age, extended model without age as covariate; IM no age, initial model without age as covariate.

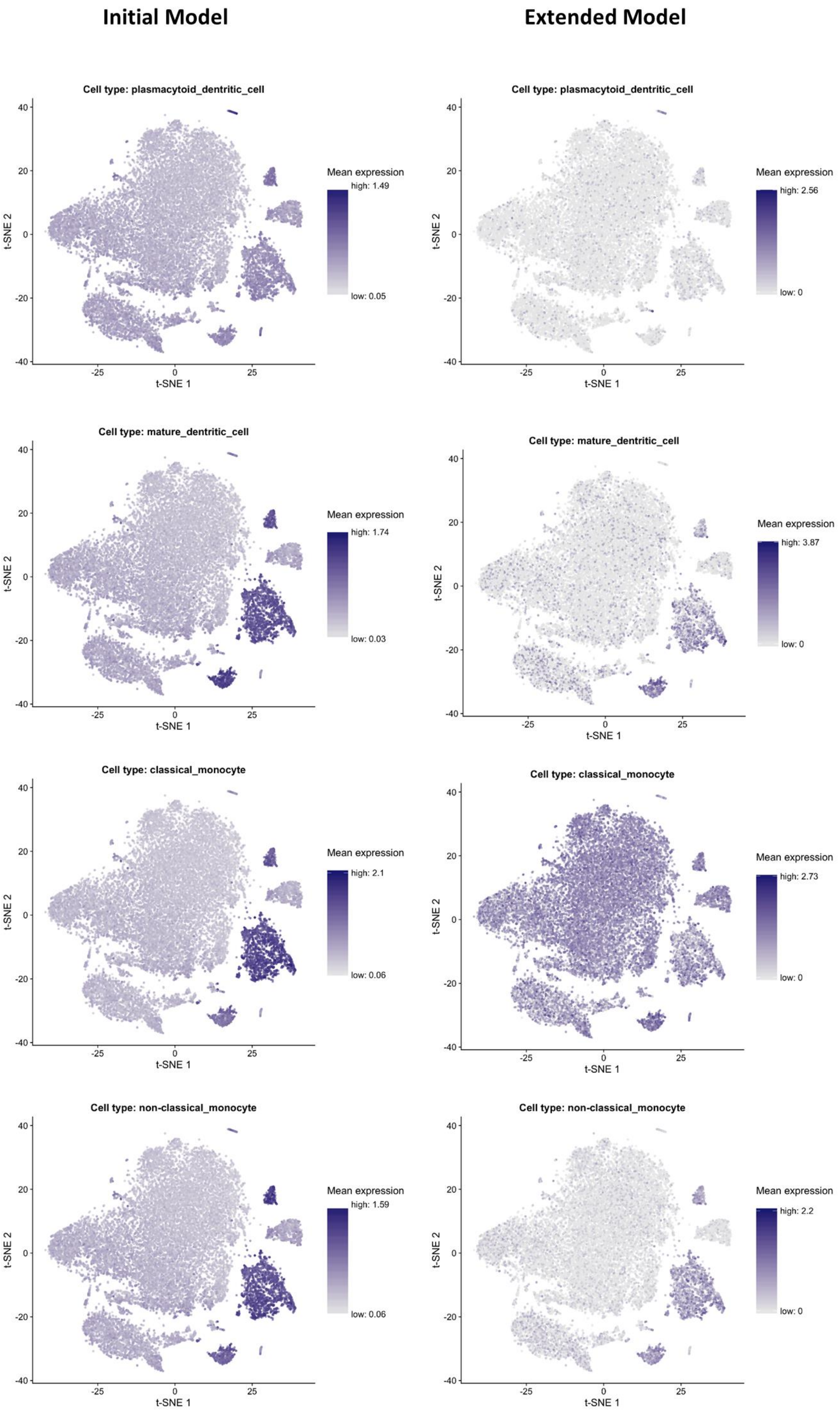


**Fig. S4.** *Continues*

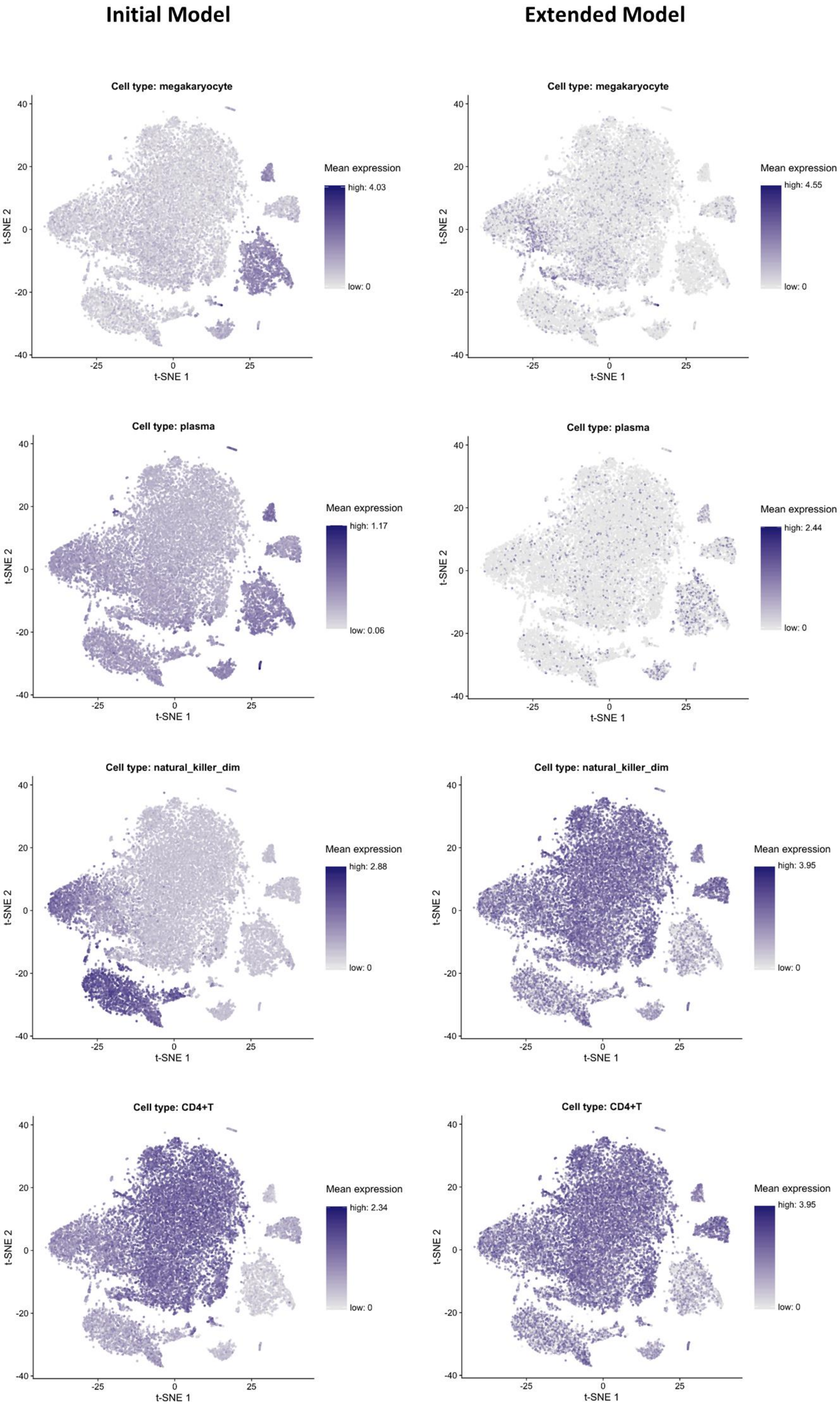


**Fig S4.** *Continues*

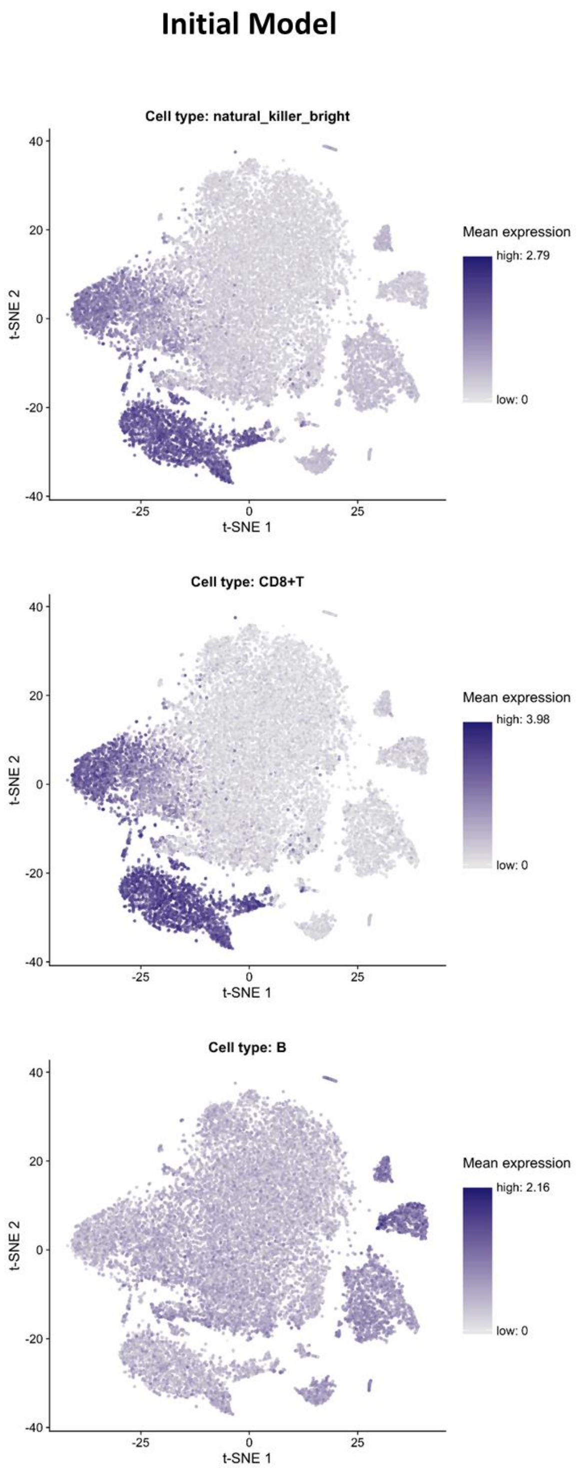


**Fig S4. scRNA-seq data-derived t-SNE plots reveal that IM-related aging genes are more likely cell type-specific marker genes.** Mean expression levels of cell type marker genes among aging-related genes identified in the Initial Model (IM, left) and in the Extended Model (EM, right) are plotted. Where applicable, IM- and EM-related intensities for same cell types plots were compared through a Wilcoxon test, always observing a P $\leq 2.2\times10^{-16}$. For details regarding cell population-specific regions, refer to [20].

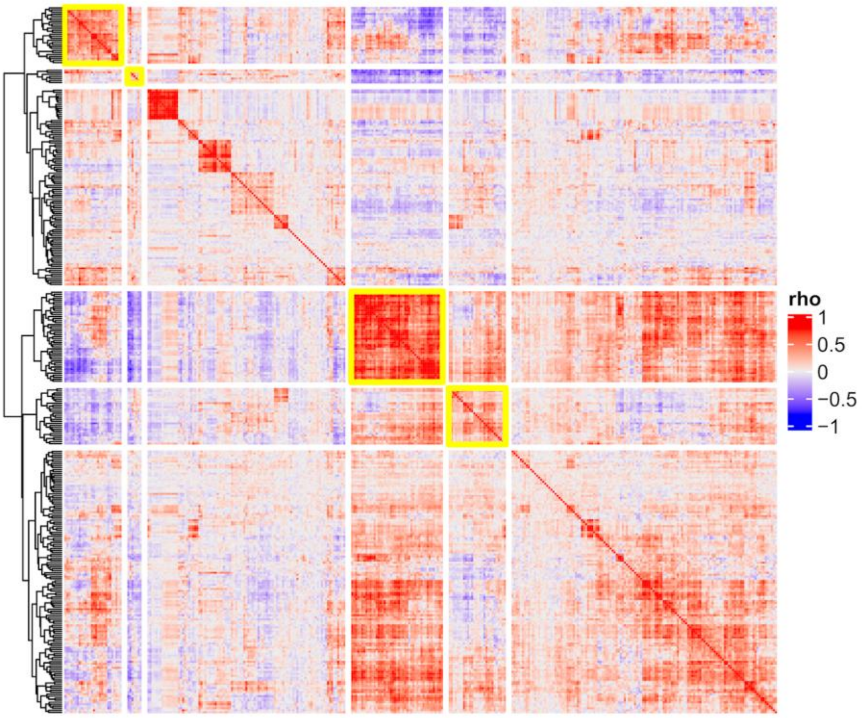


**Fig. S5. Heatmap of gene expression residuals correlations for EM downregulated aging-related genes.** Downregulated EM aging-related genes were clustered based on the correlations of gene expression residuals and highly correlating clusters were identified and highlighted with a yellow border. See *Results* section for details.

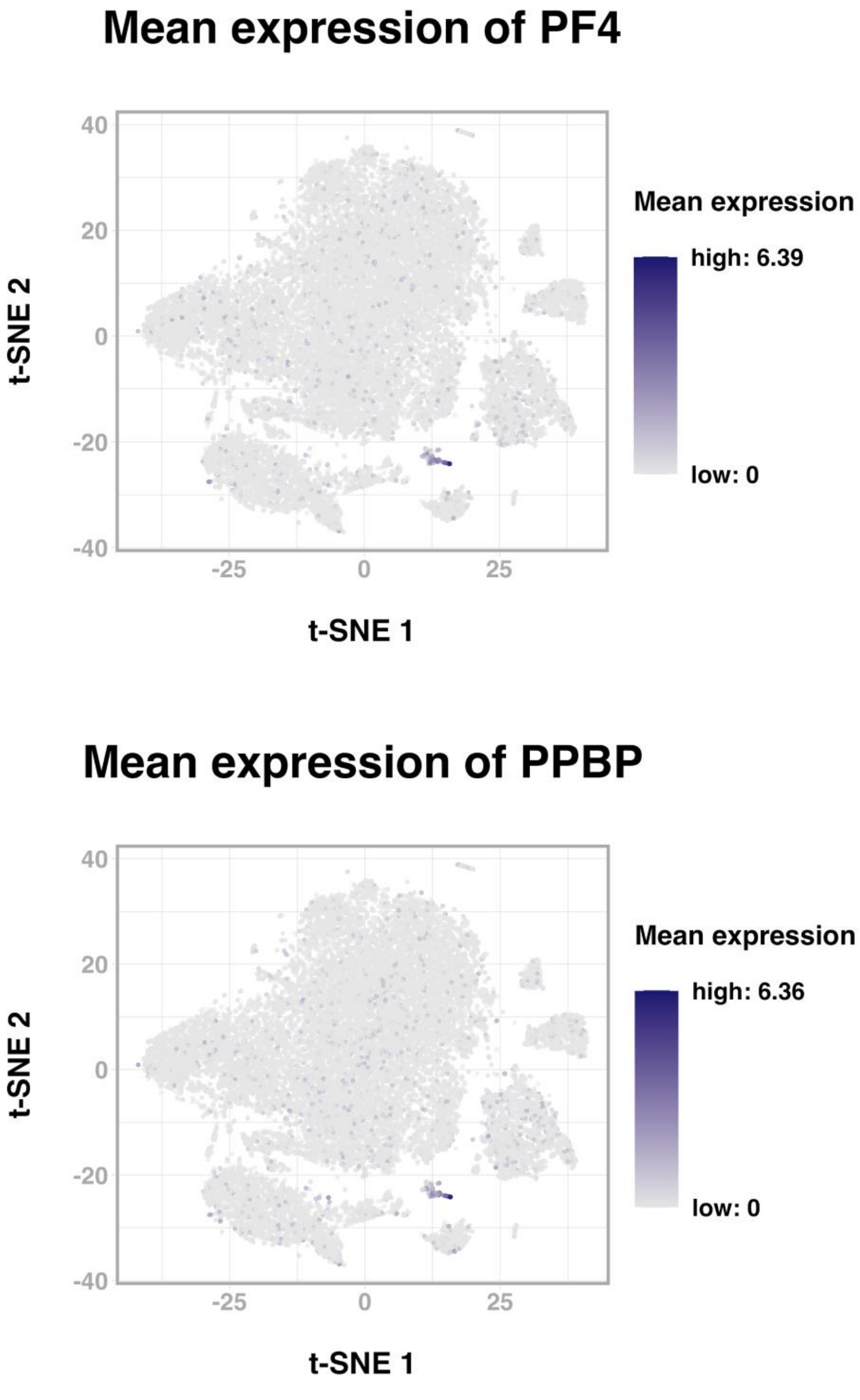


**Fig S6. scRNA-seq data-derived t-SNE plots reveal that *PF4* and *PPBP* are specifically expressed in megakaryocytes.** Mean expression levels of platelet marker genes *PF4* and *PPBP* are plotted. For details regarding cell population-specific regions, refer to [20].
